# *Hippocampal Egr1*-dependent neuronal ensembles negatively regulate motor learning

**DOI:** 10.1101/2020.11.26.399949

**Authors:** Veronica Brito, Enrica Montalban, Anika Pupak, Mercè Masana, Silvia Ginés, Jordi Alberch, Claire Martin, Jean-Antoine Girault, Albert Giralt

## Abstract

Motor skills learning is classically associated with brain regions including cerebral and cerebellar cortices and basal ganglia. Less is known about the role of the hippocampus in the acquisition and storage of motor skills. Here we show that mice receiving a long-term training in the accelerating rotarod display marked transcriptional changes in the striatum and hippocampus when compared with short-term trained mice. We identify *Egr1* as a modulator of gene expression in the hippocampus during motor learning. Using mice in which neural ensembles are permanently labeled in an *Egr1* activity-dependent fashion we identify ensembles of *Egr1*-expressing pyramidal neurons in CA1 activated in short- and long-term trained mice in the rotarod task. When *Egr1* is downregulated or these neuronal ensembles are depleted, motor learning is improved whereas their chemogenetic stimulation impairs motor learning performance. Thus, *Egr1* organizes specific CA1 neuronal ensembles during the accelerating rotarod task that limit motor learning.

## Introduction

It is now well established that the hippocampus is involved in the formation of detailed cognitive ‘maps’ of the context in which learning occurs^1^. On the other hand, the dorsal striatum is important for learning and choosing actions during procedural or motor learning^2^. However, hippocampal and striatal memory systems can operate in parallel as part of a dynamic system which optimizes behavior and the information contained in the two systems can either cooperate or compete^3^. Thus, it exists a complex and functional striatum-hippocampus interaction mostly demonstrated in human studies^4^. Supporting this functional connectivity, physical connections between the dorsal hippocampus and the ventral striatum are reported in rodents^5,6^.

In addition to the classical roles of the hippocampus in the modulation of contextual information, in the generation of cognitive maps, and in the acquisition and storage of declarative memories, this brain region also plays relevant roles in tasks associated with the striatum. For example, the hippocampus is activated during goal-directed behaviors and strategies^7,8^ and modulates contextual associations during drug-of-abuse administration^9,10^ or appetitive conditioning^11^.

In contrast, the role of the hippocampus in motor learning is poorly understood. Potential contributions of the hippocampus to motor learning and coordination have been proposed on the basis of human studies and imaging approaches^12^. Cooperative or competitive interactions between the human striatum and hippocampus seem to coordinate and synchronize during the acquisition of motor abilities^13–15^, but the underlying molecular mechanisms and the neural ensembles involved remain unknown. The use of rodent models can help to elucidate this question. Initially, although hippocampal lesion studies showed improvements in cued-learning^16^ or object recognition memory^17^, changes in motor learning tasks such as the rotarod have not been observed^18,19^. However, neuroimaging approaches have shown that the hippocampus displays higher rates of micro-structural changes in rotarod-trained mice compared to untrained controls and, accordingly, this brain region is larger in the best performers^20^. Furthermore, the accelerating rotarod task is capable to induce Fos (a marker of neural activation) labeling^21^, mTOR and cAMP-dependent protein kinase activation^22,23^, and neurogenesis^24^ in the hippocampus. These observations indicate that the hippocampus is recruited and likely to play a role in the modulation of motor skills learning.

In the present work we show how the activity in the dorsal hippocampus changes during motor learning. We also show major hippocampal transcriptional changes in long-term trained mice in the accelerating rotarod task when compared with untrained or short-term trained mice. Transcriptional profiling indicates that *Egr1* (a.k.a. *Zif-268*, *NGFI1* or *Krox24*) in the hippocampus could be a major molecular player in the regulation of motor skills learning and memory. We identify *Egr1* activity-tagged neuronal ensembles in CA1 and show that their depletion improves motor learning whereas their activation has a negative and selective impact. Thus, our results reveal the existence of hippocampal neuronal ensembles that tightly modulate the rates of motor skills learning.

## Results

### Functional interaction of striatum and hippocampus during motor learning

Hippocampal cell populations are activated during simple locomotion^25^ and during a motor learning task^12^. Less clear is whether this activation is cooperative, independent or competitive with the rest of the motor systems such as the striatum or motor cortex^3^. Thus, we first aimed to test whether the hippocampus is functionally connected with motor learning systems such as the striatum. We subjected a group of lightly anesthetized wild type mice to an fMRI scan. Seed-based analysis was performed to evaluate connectivity of the left striatum respect with the right striatum and the left and right hippocampi. We observed that the right striatum as well as both hippocampi, were positively correlated with the left striatum demonstrating that these brain regions are functionally connected (Fig. 1a-b). To better understand the hippocampal dynamics during the acquisition of a motor skill, we transduced dorsal CA1 pyramidal cells with AAV-GCaMP6f and we placed a fiber-optic probe to monitor pyramidal neurons activity in CA1during the accelerating rotarod task (Fig. 1c-d). We observed two interesting phenomena. First, during the initial trials of the accelerating rotarod task the Ca^2 +^-dependent signal dynamics were irregular and heterogenous whereas at the end of the task the signal progressively decreased and stabilized (Fig. 1e-f). Second, we observed Ca^2 +^-dependent peaks just before the mice fell from the rotating rod. These peaks were detected only during the first 1-2 days of training but decreased at the end of the task, especially on days 4-5 (Fig. 1g-h). Altogether, these results suggest that the pyramidal cells of the CA1 progressively reduce their activity across trials in the rotarod task and, that they have a predictive value for falls at the beginning of the training.

**Figure 1.**
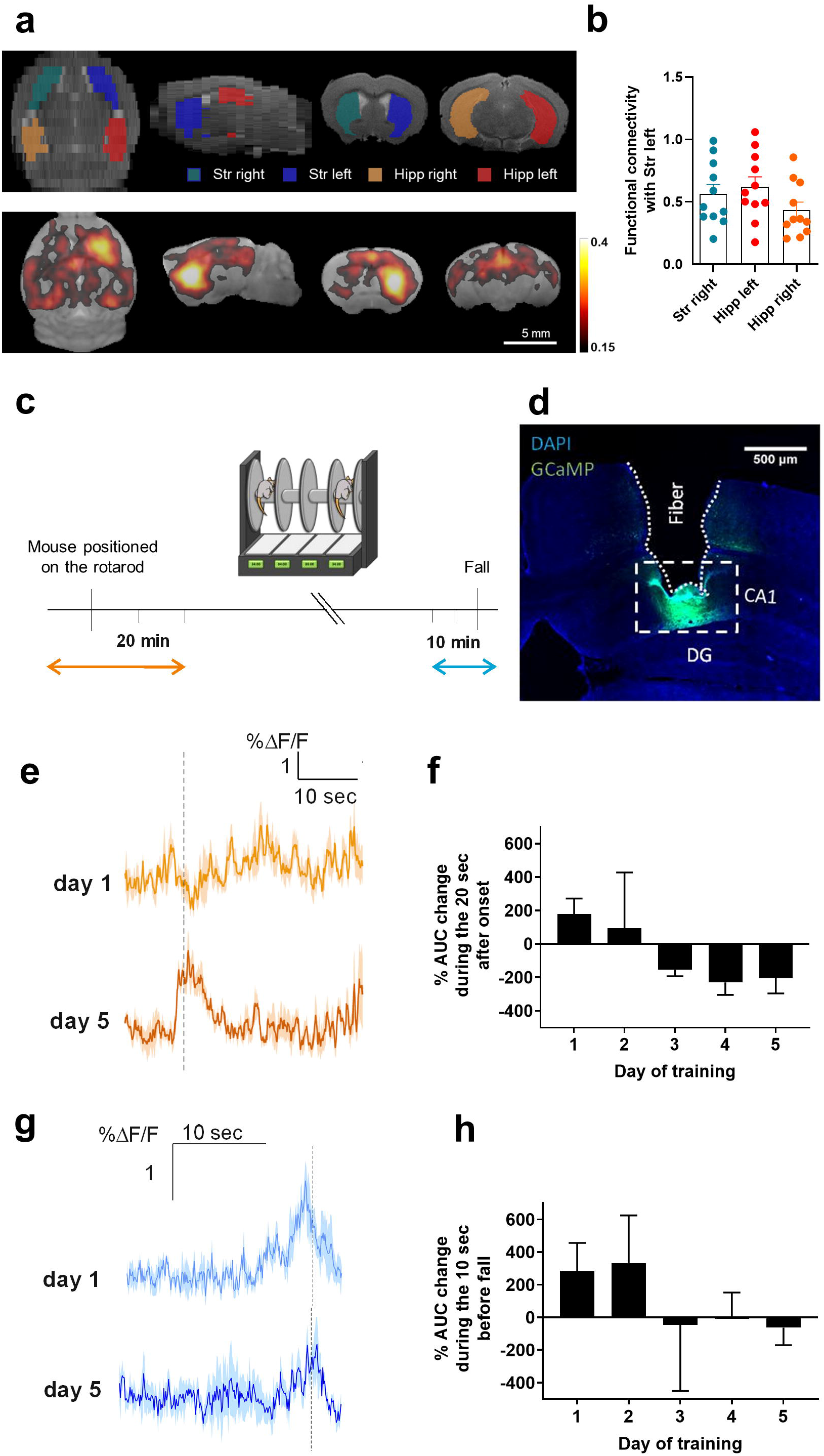
Basal striatal-hippocampal functional connectivity and calcium dynamics in CA1 during the accelerating rotarod task. (**a**) Average seed-based BOLD correlation maps from left striatum in wild type mice. A: Anterior, P: Posterior, L: Left, R: Right, S: Superior, I: Inferior. (**b**) Mean correlation value of striatum (Str) with hippocampus (Hipp) in both hemispheres. Data are scatter plots with mean ± SEM (n = 11 mice). (**c**) Experimental design of fiber photometry. Analysis was performed during the time windows indicated by orange (beginning) and blue (end of trial) arrows. (**d**) Viral expression of AAV-GCaMP6f and placement of the fiber-optic probe in CA1. DG, dentate gyrus. (**e**) Representative Ca^2 +^ traces from a mouse during the beginning of trials (dashed lines indicate the moment the mouse was placed in the rotarod) on the first and last training sessions (days 1 and 5). Data are percent change in fluorescence over the mean fluorescence (%∆F/F). (**f**) Quantification of the percentage of change in the area under the curve (AUC) between □0, 10 sec] and [10, 20 sec] after the animal is placed in the rotarod. (**g**) Representative Ca^2 +^ traces from a mouse during the end of trials (when the animal falls, dashed line) on days 1 and 5. Data the percent change in fluorescence over the mean fluorescence (%∆F/F). (**I**) Quantification of the percentage of change in AUC between [-10, −5 sec] and [−5, 0 sec] before the animal falls from the rotarod.

### Differential gene regulation in striatum and hippocampus during motor learning

To characterize potential changes occurring in the hippocampus during the acquisition of a motor skill, we looked for global transcriptional alterations. We subjected two groups of mice to the accelerating rotarod task (Fig. 2a-b). The first group (short-term trained or STT) was trained just one day in the task to assess the initial phase of motor learning, and a second group was trained for five days to acquire well-learned motor skill (long-term trained or LTT). Both groups were compared with mice that were not trained but were placed in the apparatus as a control (non-trained or NT). Twenty-four hours after the last day of exposure to the rotarod or training, the dissected dorsal hippocampus (DHipp) and dorsal striatum (DStr) of the mice from the three groups were subjected to deep sequencing analysis (RNAseq). We first compared the genes differentially expressed (DEGs) between DStr and DHipp using a paired-sample analysis of NT samples. In the non-trained condition, we found that around 30% of all detected genes (4,927 of 16,915 genes, Supplementary table 1) were differentially expressed between DStr and DHipp (Adj p-value <0.01; log2FC > 0.3 or < −0.3; Fig. 2c). We then examined the differences between regions in the short and long training conditions (Supplementary table 1) and a Venn diagram was used to classify DEGs between DHipp and DStr found in all three datasets (NT, STT and LTT). Most DEGs (3921) showed similar expression signature between all conditions, with 56% showing increased expression in DHipp and the rest in DStr. In addition, we observed that several genes were differentially expressed in only one or two of the three conditions (Fig. 2d). To assess the functional profile of DEGs enriched in the DStr and DHipp we used enrichment analysis by mapping to KEGG database. We found that DEGs related to dopamine receptor signaling was significantly enriched in DStr (Fig. 2e), whereas those associated with glutamate signaling were prevalent in DHipp (Fig. 2f).

**Figure 2.**
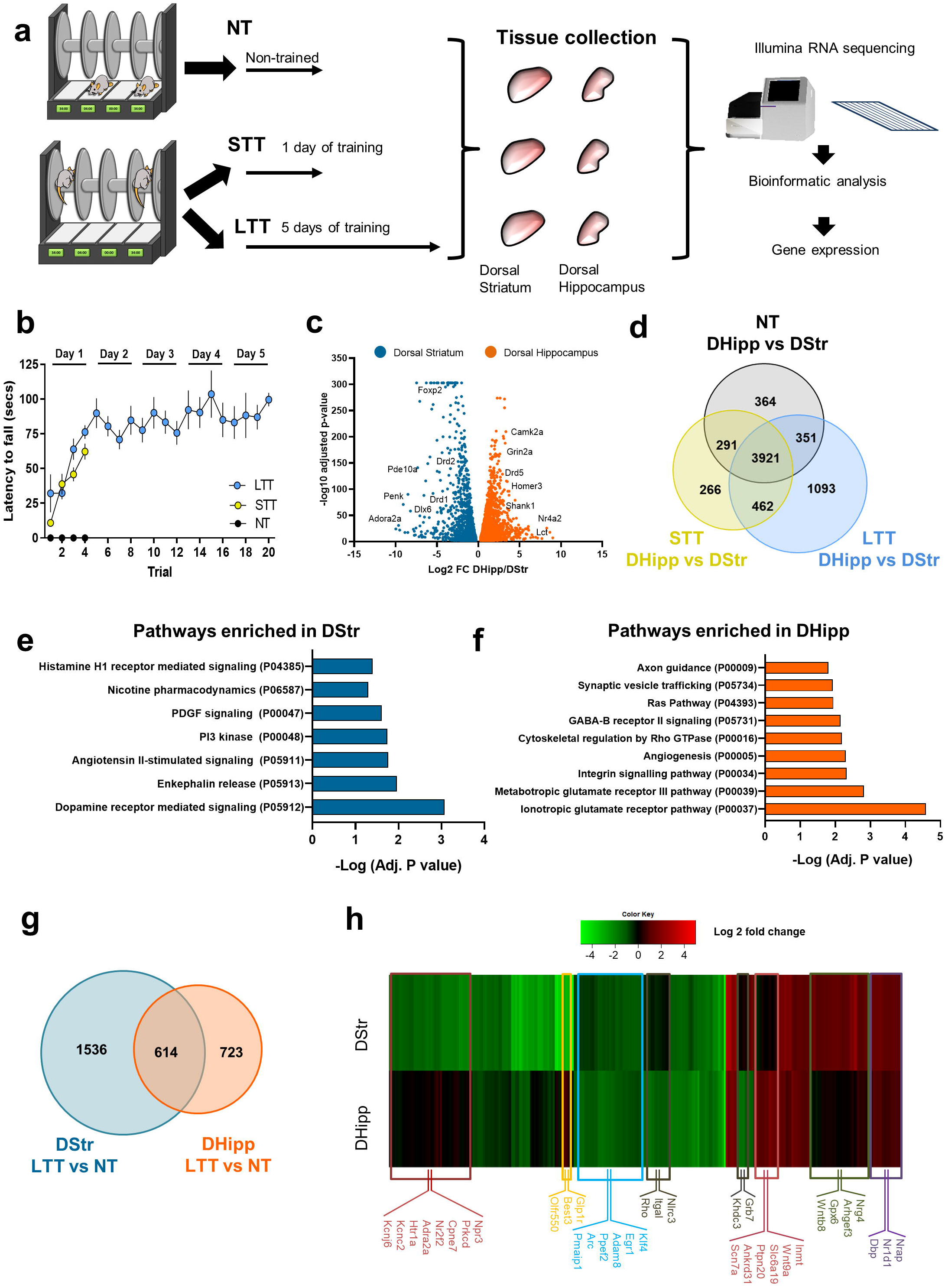
Striatal and hippocampal gene expression profile following the accelerating rotarod task. (**a**) Schematic of experimental design. WT mice were trained on a rotarod. Three groups were evaluated (n = 8 per group): non-trained (NT), short-term trained (STT), and long-term trained mice (LTT). The transcriptome of the 5 best performers in each group was analyzed by RNA sequencing 24 h after the last trial. (**b**) Latency to fall in the accelerating rotarod task in NT, STT, and LTT mice. Data are means ± SEM. (**c**) Volcano plots showing significant mRNA differential expression genes between DStr (blue) and DHipp samples (orange) in non-trained mice (Adj p-value <0.01; log2FC > 0.3 or < −0.3). The name of representative mRNAs is indicated. (**d**) Venn diagram of data in Supplementary table 1 showing the number of mRNAs differentially expressed in DHipp vs DStr samples in NT (grey), SST (yellow) and LTT (light blue) mice. (**e, f**) Pathway analysis for transcriptomic enrichment in DStr (**e**) vs. DHipp (**f**) by using Panther analysis including all significant differentially expressed genes (FDR< 0.05) in LTT (**g**) Venn diagram of data in Supplementary table 3 showing the number of mRNAs differentially expressed in LTT vs NT samples in the DStr (blue) and DHipp (orange). (**h**) Heatmap depicting the top differentially expressed genes in either regions after LTT (padj<0.01; log2FC= ± 1). Colors of heatmap represent log2FC. Names of representative mRNA are indicated.

Next, we assessed the overall transcriptional changes in response to training comparing LTT vs NT and STT vs NT in the DStr and in the DHipp (Fig. 2g and Supplementary table 3, Adj p-value <0.05; log2fold change > 0.3 or < −0.3). The comparison of the transcriptional profile between STT and NT showed significant changes in DHipp or DStr for only few genes (Supplementary Table 3). In contrast, the comparison between NT and LTT revealed that a large number of genes were down- and up-regulated in DStr (1132 and 1018 respectively) and in DHipp (797 and 540, respectively). Remarkably, although many LTT vs NT differences were common to DStr and DHipp, a unique gene expression profile was observed in each region in response to LTT as shown by the Venn diagram in figure 2g. The most significant changes in gene expression driven by motor learning training in DHipp and Dstr are represented in the heat map in figure 2h. Hierarchical clustering of gene expression changes in LTT vs NT revealed major genes clusters showing 1) genes regulated in an opposite manner in the two regions (i.e. upregulated in DHipp and downregulated in DStr or vice versa), 2) genes with similar changes in expression, and 3) genes with changes in expression unique to either one region or the other (Fig. 2h).

To identify potential transcription factors specifically regulating DEGs, we searched for transcription factor binding motifs in the genes differentially expressed between the DHipp and DStr in the three conditions (see data sets in Supplementary table 1). The analysis revealed that promoter regions of DEGs between DHipp and DStr were highly enriched for *SP1*, *Egr1*, *SMAD4*, *TEAD2* binding motifs in NT, ST and LTT (Fig. 3a and 3d, Supplementary table 4). Interestingly, among these transcription factors, only *Egr1* gene expression levels were significantly changed after LTT (Fig 3b and 3e) in both, the DStr and DHipp. This result was confirmed by qPCR (Fig 3c and 3f). Furthermore, we observed that 7% of the hippocampal genes whose expression changed in response to LTT (comparison LTT vs NT) are regulated by *Egr1* (62 are downregulated and 32 upregulated, Supplementary Table 5). Interestingly, using the SynGo platform, we identify in this gene data set, 11 genes mapping with processes in synapse assembly, organization and function, and regulation of postsynaptic membrane neurotransmitter receptor levels (Supplementary Table 6). Taken together these data indicate that during the progressive learning of a motor skill, long-term transcriptional changes associated with *Egr1* activity occur in both hippocampus and striatum and are presumably associated with its acquisition and/or maintenance.

**Figure 3.**
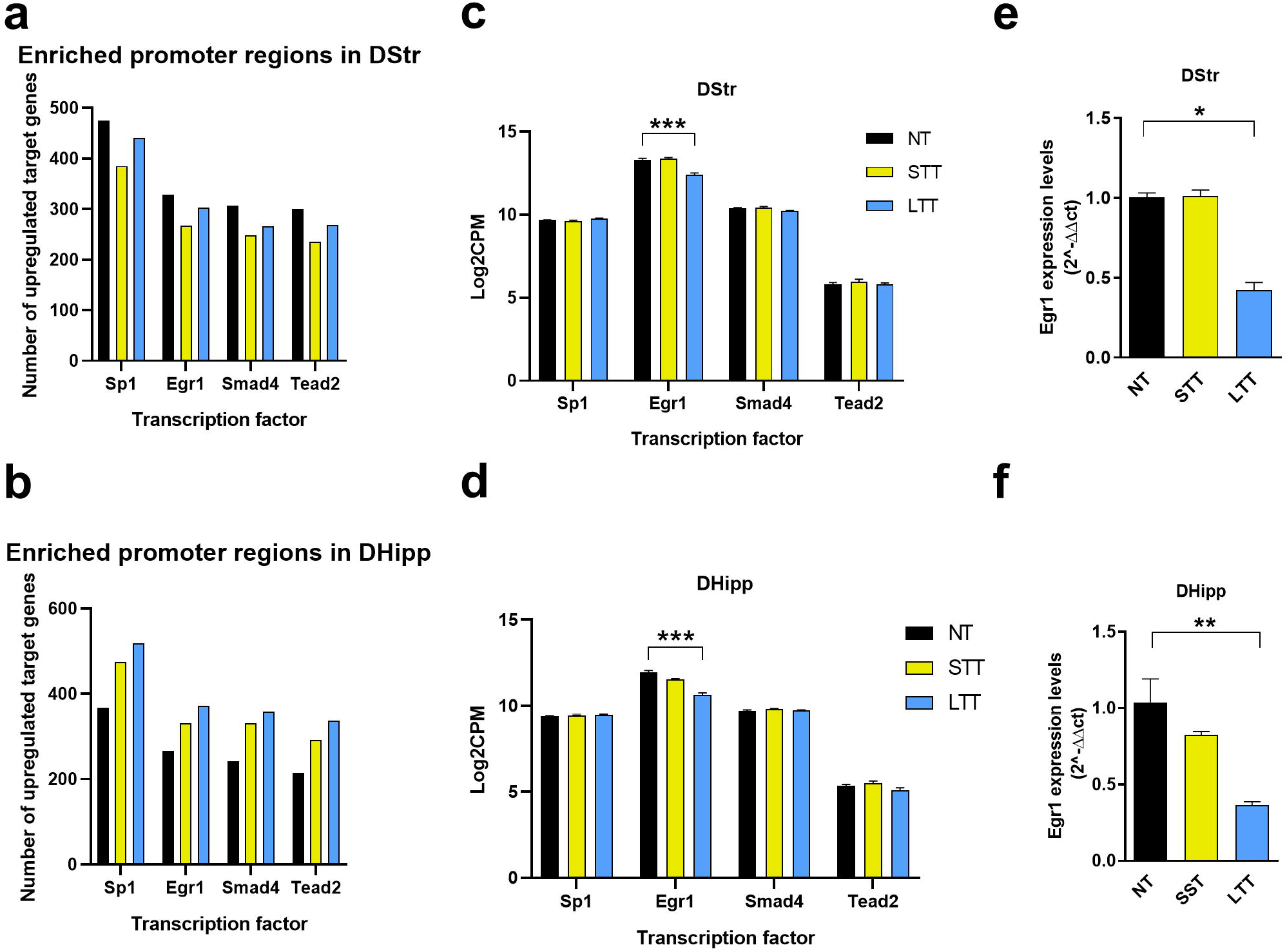
Characterization of transcription factor binding sites motifs in genes differentially regulated in the dorsal striatum and dorsal hippocampus following accelerating rotarod training. (**a, b**) Binding motifs detected at gene promoters using Enrich Tool through scanning the TRANSFAC and JASPAR databases. Plots show the 4 most significant transcription factors with the number of predicted target genes that are upregulated in the DStr (**a**) or in the DHipp (**b**) in NT, STT, and LTT samples (FDR < 0.01). (**c, d**) Gene expression levels of *Sp1*, *Egr1*, *Smad4* and *Tead2*transcription factors in the DSt (**c**) and DHipp (**d**) are represented as log2 counts (*p<0.001 NT vs LTT). (**e, f**) *Egr1* gene product levels measured by quantitative RT-PCR in DStr (**e**) and DHipp (**f**) in NT, ST and LTT mice samples. Scatter plot in **e** and **f** with means ± SEM (n = 5 per group). Kruskal-Wallis test identified general significant changes between groups in **e**(K-W statistic = 9.42; *p* < 0.0025) and **f**(K-W statistic = 10.24; *p* < 0.0006). Dunn’s *post hoc* analysis (* *p* < 0.05, ** *p* < 0.01, *** p<0.001) compared with the NT group.

### *Egr1*-dependent engrams are activated in CA1 during motor learning

From the transcriptomic profiling experiments, we identified the *Egr1* gene product as a potential molecule that could play a significant role in the hippocampus during the neural plastic changes occurring in different phases of motor learning in the rotarod task. To explore the role of *Egr1* in motor learning we used recently generated *Egr1*-CreER^T2^ BAC transgenic mice^26^ (Fig. 3a). These mice express the Cre recombinase fused to modified estrogen receptor (ER^T2^) under the control of the *Egr1* promoter^27^. In the fusion protein, Cre activity is induced by exogenous treatment with 4-hydroxytamoxifen (4-HT). To identify which neural cells are activated during different phases of motor learning, we crossed *Egr1*-CreER^T2^ mice with R26^RCE^ mice, a reporter line in which EGFP expression requires recombination by Cre (Fig. 4a). In double transgenic *Egr1*-CreER^T2^ x R26^RCE^ mice, cells in which *Egr1* is induced in the presence of 4-HT become permanently labeled with EGFP. We subjected *Egr1*-CreER^T2^ x R26^RCE^ mice to the accelerating rotarod task using same three groups as above (NT, STT, and LTT, see Fig 2a). Each group received a single injection of 4-HT 1 h before the last session of training in the task (Fig. 4b). Three days (72 h) after the 4-HT injection, brains were processed for immunofluorescence staining and confocal imaging. We counted GFP-positive neural cells in several brain regions: dorsal striatum, ventral striatum, layers 2/3, 4, 5, and 6 of the motor cortex area 1, and the CA1 and CA3 of the dorsal hippocampus (Supplementary fig. 1). The density of GFP-positive neural cells was increased in CA1 in both STT and LTT groups compared with NT group (Fig. 4c-d) whereas it was increased only in the STT group in layers 5 and 6 of the motor cortex area 1 (Supplementary fig. 2a-b, Fig. 4d) and in the dorsal striatum (Supplementary fig. 2a and c, Fig. 4d). These results indicated a more sustained *Egr1* activation in a specific group of neural cells in CA1 compared with the dorsal striatum and the motor cortex. We then characterized these GFP-positive neural cells. Co-localization studies revealed that most GFP-positive cells in the dorsal striatum (Supplementary fig. 3a) were DARPP-32-positive striatal projection neurons (SPNs) (Fig. 4f). Most GFP-positive cells in the motor cortex area 1 and CA1 (Supplementary fig 3b-c) were NeuN positive, indicating their neuronal identity (Fig. 4f). Finally, most GFP-positive cells in CA1 were NeuN and MAP2-positive (Fig. 4e-f) and parvalbumin-negative (Supplementary fig. 3c and fig. 3f) indicating they were pyramidal neurons (Fig. 4e-f).

**Figure 4.**
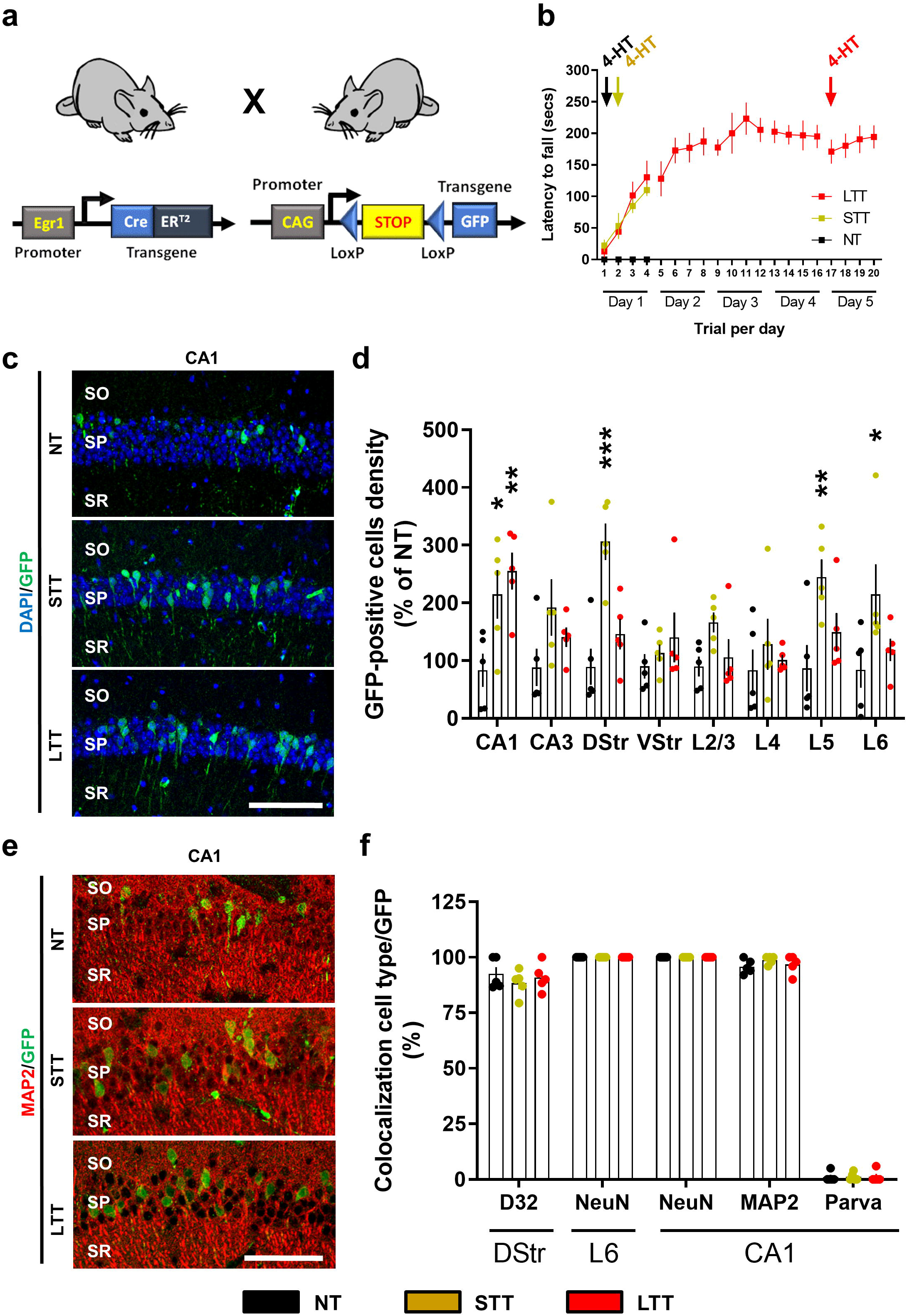
*Egr1*-dependent neural ensembles during different phases of the rotarod task training. (**a**) Schematic representation of mutant mice used. (**b**) We subjected these mice to three different conditions in an accelerating rotarod task (as in **Fig. 2a**): NT, STT, and LTT. All the mice (n = 5 mice per group) received an injection of 4-HT 1 h before the rotarod training session (arrows) on the last day of training. (**c**) Representative images of *Egr1*-dependent activation of neural cells (GFP-positive, green) co-stained with DAPI (blue) in dorsal CA1. (**d**) Pseudo-stereological quantification of GFP-positive neural cells density per area in NT, STT and LTT groups of mice (n = 5 mice per group). Scatter plot with means ± SEM. Two-way ANOVA identified general significant changes between groups (F_(2, 88)_ = 16.9; *p* < 0.0001). Dunnett’s *post hoc* analysis (* *p* < 0.05, ** *p* < 0.01, *** p<0.001) compared with the NT group. (**e**) Representative images of *Egr1*-dependent activation of neural cells (GFP-positive, green) co-stained with MAP2 (red) in CA1. (**f**) Quantification of the percentage of GFP-positive neural cells that co-localizes with various neural markers (in red) in the brain regions which displayed significant differences in **d**. DARPP-32 (D32) and parvalbumin (Parva). No significant changes were identified between groups (n = 5 mice per group) regarding the neural type analyzed. DStr: dorsal striatum; VStr: ventral striatum; L2/3, L4, L5 and L6: motor cortex layers 2-6. CA1-CA3: *cornu ammonis*. SO: *stratum oriens*, SP: *stratum pyramidale*, SR: *stratum radiatum*. Scale bar in C and D, 100 μm.

### *Egr1* knockdown in CA1 potentiates motor performance

To test whether *Egr1* levels in the dorsal CA1 control the acquisition and/or maintenance of the motor skills required for the accelerating rotarod task we transduced the CA1 of wild type (WT) mice with an adeno-associated virus (AAV) expressing a shRNA against the *Egr1* transcript (shRNA-*Egr1* group, fig. 5a) or a control shRNA (scramble group) (Fig. 5b). After three weeks of viral transduction, we observed a significant reduction of *Egr1* immuno-reactivity in the pyramidal cells of the dorsal CA1 in shRNA-*Egr1* mice compared with the scramble group (Fig. 5c-d). To test the effects of down-regulating *Egr1* in the CA1 we subjected the shRNA-*Egr1* and scramble groups of mice to the accelerating rotarod task (Fig. 5e). Scramble mice progressively learned the task and reached a plateau of performance from the second day of training on, whereas, unexpectedly, the shRNA-*Egr1* mice kept improving their performance until the 4^th^ day of training (Fig. 5e). This effect was accompanied by significant increased latencies to fall from the rotarod in shRNA-*Egr1* mice compared with the scramble mice. These results provide evidence that *Egr1* down-regulation in the pyramidal neurons of the CA1 improves the accelerating rotarod performance.

**Figure 5.**
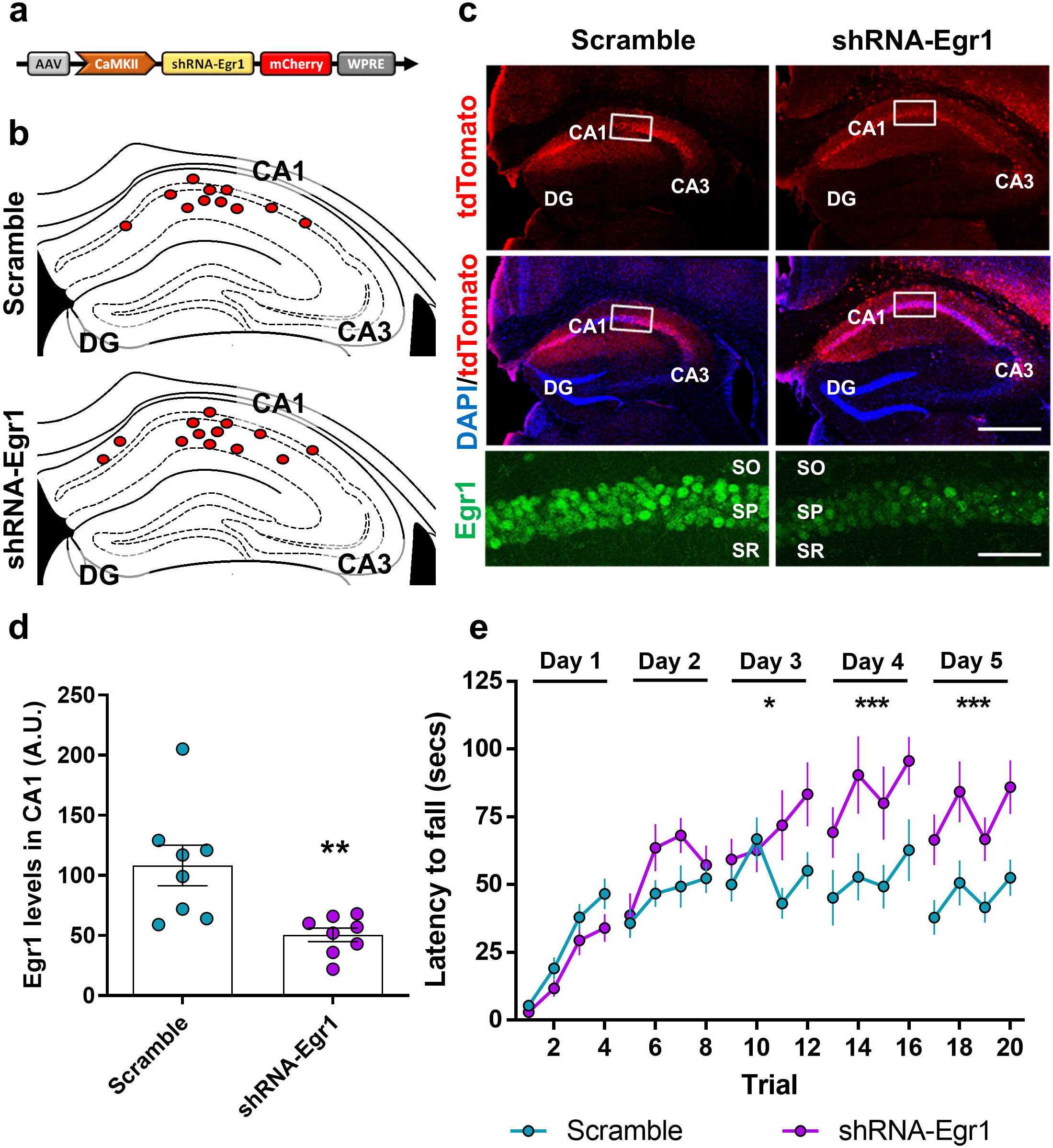
Effects of *Egr1* down-regulation in CA1 on accelerating rotarod training. (**a**) Schematic illustration of the transduced adeno-associated virus (AAV-pCaMKII-shRNA-*Egr1*-mCherry) in WT mice. (**b**) Schematic location of the injection site centers in dorsal CA1 for all mice used in this experiment (only left hemisphere is shown). (**c**) Representative images of hippocampi transduced with control AAV (AAV-pCaMKII-Scramble-mCherry, Scramble, left) or with the experimental AAV (AAV-pCaMKII-shRNA-*Egr1*-mCherry, shRNA-*Egr1*, right). Triple staining showing all cells (DAPI, blue), transduced cells (mCherry, red), and *Egr1*-positive cells (Inset, green) for each group of mice. Scale bars, 300 μm, inset: 60 μm. (**d**) Quantification of *Egr1* optical density (arbitrary units) in the pyramidal cell layer of CA1. Data are scatter plots (one point per mouse, average from two slices per mouse) and means ± SEM Mann-Whitney t test, sum of ranks A, 95, B, 41, Mann-Whitney U, 5, *p* = 0.003, **. N = 8 mice per group. (**e**) Three weeks after viral transduction mice were subjected to the accelerating rotarod task for five days (4 trials per day). N = 11 per group. Data are means ± SEM and analyzed by two-way ANOVA. A significant difference was detected between groups on day 3, F_(1, 80)_ = 6.30, *p* = 0.0141), day 4, F_(1, 80)_= 16.91, *p* < 0.0001), and day 5, F_(1, 80)_ = 26.03, *p* < 0.0001. DG: Dentate gyrus, CA1: *cornu ammonis* 1, CA3: *cornu ammonis* 3, SO: *stratum oriens*, SP: *stratum pyramidale*, SR: *stratum radiatum*.

### Depletion of CA1 *Egr1*-dependent engrams enhances motor performance

We observed *Egr1*-dependent neuronal ensembles formation in CA1 during the acquisition of motor skills in the accelerating rotarod task and we showed that a global down-regulation of *Egr1* in the dorsal CA1 enhances the performance in this task. We therefore tested the consequences of depleting CA1 *Egr1*-dependent neuronal ensembles on motor learning to evaluate their contribution. We used the double mutant *Egr1*-CreER^T2^ x R26^RCE^ mice (Fig. 6a). These mice were transduced bilaterally in the dorsal CA1 with vehicle or AAV-flex-taCasp3-TEVp (Fig. 6b). Three weeks later, all mice were subjected to the accelerating rotarod task (Fig. 6c) receiving an i.p. injection of 4-HT 1 h prior the training session. This injection was administered at days 1 and 2 of the task. With this design, specific *Egr1*-dependent CreER^T2^ induction in activated CA1 pyramidal cells of the CA1 would induce Caspase-3 expression in the presence of 4-HT resulting in cell death of the hippocampal components of *Egr1*-dependent neuronal ensembles. Mice transduced with AAV-flex-taCasp3-TEVp (Casp3 group) in CA1 displayed higher latencies to fall on days 3 and 5 (on day 4 there was just a trend) compared with control mice (Vehicle group) (Fig. 6c). Brains from these mice were examined to verify the depletion of the *Egr1*-dependent neuronal ensembles (Fig. 6d-e). As expected, mice infused with vehicle in CA1 showed an increase in GFP-positive cells in the pyramidal layer of CA1 after rotarod training when compared with NT mice (as previously shown in fig. 4). In contrast, the density of GFP-positive cells in the dorsal CA1 was dramatically reduced in rotarod-trained mice that were transduced with Casp3 (Fig. 6d-e). This latter result shows that the *Egr1*-dependent neuronal ensembles were depleted. This depletion was accompanied by a better performance in the rotarod task of Casp3-transduced mice as compared with the vehicle-injected mice that received the same training and 4-HT treatment.

**Figure 6.**
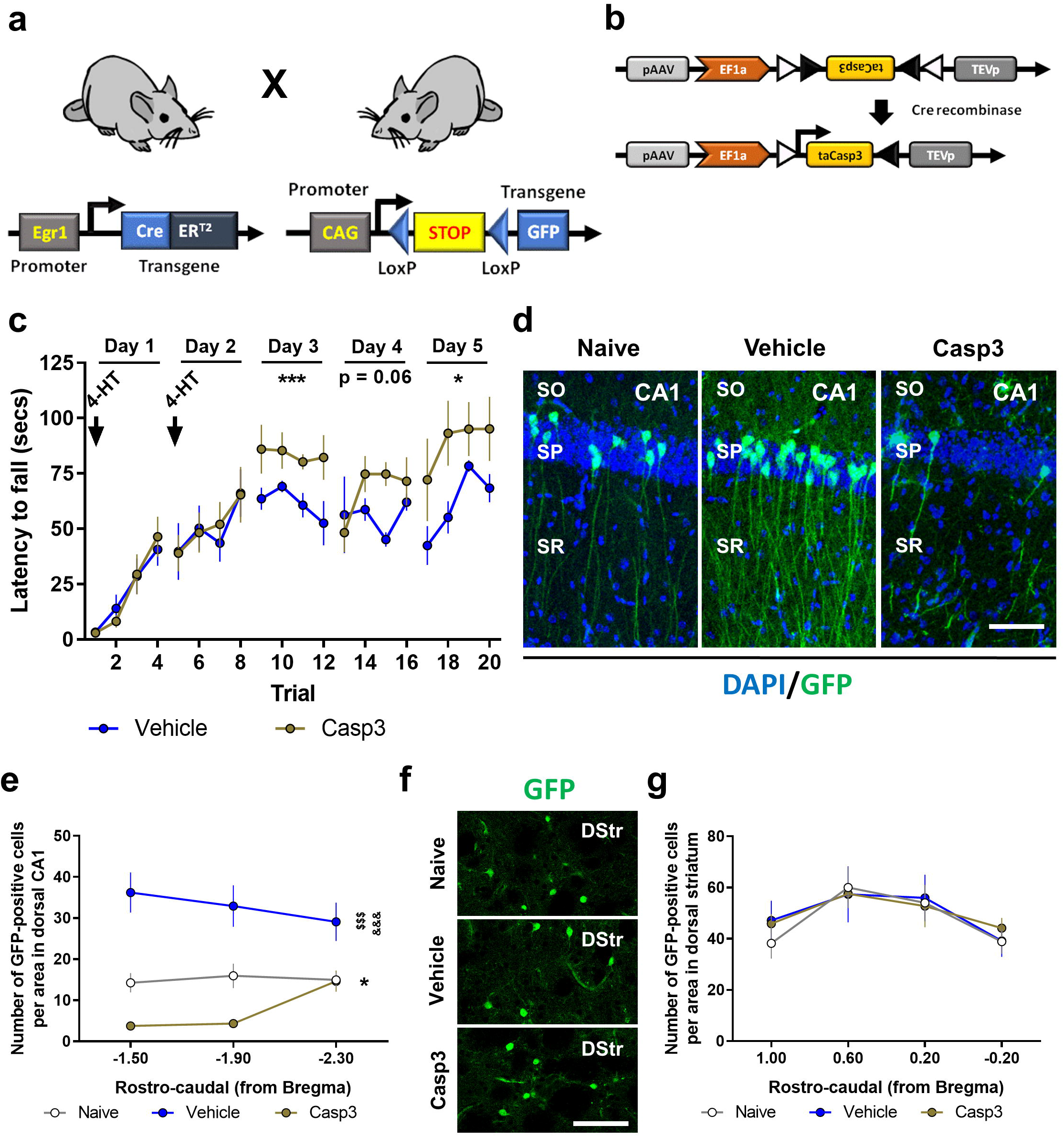
Depletion of the CA1 *Egr1*-dependent neuronal ensembles during the rotarod task. (**a**) Schematic representation of the double mutant mice used in this experiment. (**b**) Schematic representation of the AAV-flex-taCasp3-TEVp vector. Mice were bilaterally injected in CA1 with AAV-flex-taCasp3-TEVp (Casp3, n = 8) or vehicle (Vehicle, n = 7). A group of non-trained mice (Naive, n = 5) was also used. (**c**) All three groups received a single injection 4-HT 1 h prior to the rotarod training (arrows). Data are means ± SEM, analyzed by two-way ANOVA. A significant difference was detected between groups on day 3, F_(1, 52)_ = 15.78, *p* = 0.0002, and day 5, F_(1, 52)_ = 10.05, *p* = 0.0025. (**d**) Representative images of *Egr1*-dependent activation of neural cells (GFP-positive, green) co-stained with DAPI (blue) in the dorsal CA1 of the hippocampus in the three groups. Scale bar, x60 μm. (**e**) Quantification of GFP-positive neural cells density per area in the three groups. Data are means ± SEM. Two-way ANOVA identified general significant changes between groups, F_(2, 51)_ = 49.27, *p* < 0.0001. Tukey’s *post hoc* analysis indicated that both, Naive (*p* < 0.05) and Casp3 (*p* < 0.001) showed significantly different GFP-positive cell density compared with Vehicle mice. **\***: Naive vs Vehicle groups; ^**$**^: Casp3 vs Naive; ^**&**^: Casp3 vs Vehicle. (**f**) Representative images of *Egr1*-dependent activation of neural cells (GFP-positive, green) in the dorsal striatum (DStr) from the three groups. (**g**) No significant differences were detected between groups. SO: *stratum oriens*, SP: *stratum pyramidale*, SR: *stratum radiatum*.

We also evaluated whether the improvement of Casp3 mice in the rotarod task could be accompanied by a higher activation of neuronal ensembles in the striatum as a consequence of the *Egr1*-engram depletion in CA1. To do so, in the same mice, we analyzed the density of GFP-positive cells in the dorsal striatum (Fig. 6e-f). There was no change in the number of *Egr1*-dependent neuronal ensembles in the dorsal striatum (Fig. 6e-f). Taken together, our results suggest an important role of CA1 *Egr1*-activated pyramidal cells in the modulation of motor skills during an accelerating rotarod task.

### Chemogenetic activation of CA1 *Egr1*-dependent engrams impairs motor performance

To further characterize the role of this *Egr1*-dependent neuronal ensemble in CA1 induced by motor learning, we used an opposite strategy, aiming to activate it using *designer receptors exclusively activated by designer drugs* (DREADDs) technology. We transduced the dorsal CA1 of *Egr1*-CreER^T2^ mice with an AAV expressing the activator DREADD hM3D(Gq) using a FLEX switch vector (Fig. 7a-b). Three weeks after AAV injection, the mice were first subjected to the accelerating rotarod task (Fig. 7c). On days 1 and 2 of rotarod training, all mice received an injection of 4-HT to induce Cre-mediated recombination in the *Egr1*-dependent neuronal ensembles. On days 4, 5 and 13 these *Egr1*-dependent neuronal ensembles were activated using clozapine-N-oxide (CNO). The same mice were also tested in the open field (day 14) and novel object location (days 15 and 16, see below). At the end of the study, to test the efficacy of hM3D(Gq) receptor stimulation by measuring cFos induction, mice were treated with CNO or vehicle 2 h before sacrifice and histological analysis (Fig. 7c). Post-mortem histology indicated that CA1 was well targeted with the vector expressing the hM3D(Gq) receptor (Fig. 7d-e). Moreover, CNO induced a robust up-regulation of cFos immunoreactivity in the transduced pyramidal cells in CA1 as compared with non-transduced cells or transduced cells from mice treated with vehicle (Fig. 7f-g).

**Figure 7.**
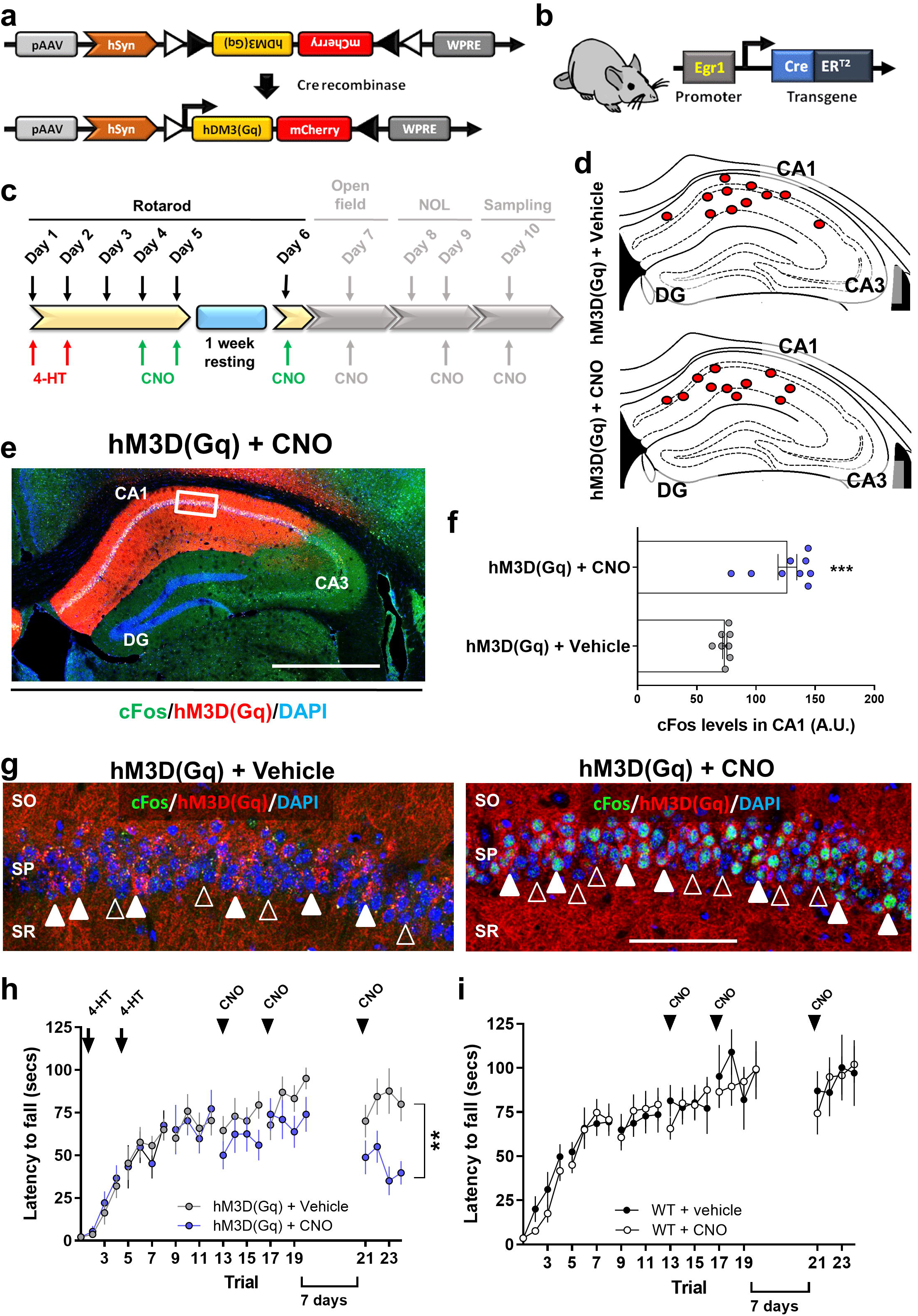
Effects of chemogenetic activation of the *Egr1*-dependent CA1 engram on the accelerating rotarod task. (**a**) Design of the AAV to express hM3D(Gq) only in cells with Cre recombinase activity. (**b**) *Egr1*-CreER^T2^ mutant mice used. (**c**) Scheme depicting the behavioral characterization in the present figure (colored). (**d**) Schematic location of the center of the injection site for each mouse (only left hemisphere is shown). (**e**) Representative image of a transduced hippocampus. (**f**) Quantification of cFos immunofluorescence intensity in the CA1 pyramidal cell layer of transduced mice and treated with 4-HT and Vehicle (n = 8) or CNO (n = 9). Means ± SEM Mann-Whitney t test (sum of ranks A, 36, B, 117, U = 0, *p* < 0.0001). (**g**) Representative images of CA1 in mice from **f**. Empty arrowheads indicate non-transduced cells, white arrowheads transduced cells. (**h**) Accelerating rotarod task in transduced mice. Arrows indicate treatment with 4-HT and arrowheads treatment with CNO. Data are means ± SEM. Two-way ANOVA identified significant differences between groups only on day 6 of training (F_(1, 76)_ = 26,41, *p* < 0.0001). (**i**) Two independent groups of WT mice treated with vehicle (n = 10) or CNO (3 mg/kg, n = 11) were subjected to the accelerating rotarod task with the same design as in **h**. Data are means ± SEM Scale bar in **e**, 300 μm, in **g**, 60 microns. DG: *dentate gyrus*, CA1: *cornu amonis* 1, CA3: *cornu amonis* 3, SO: *stratum oriens*, SP: *stratum pyramidale*, SR: *stratum radiatum*.

Concerning the rotarod performance, *Egr1*-CreER^T2^ mice transduced with hM3D(Gq) and injected with 4-HT on days 1 and 2, progressively learned the task on days 1, 2, and 3 (Fig. 7h). On days 4 and 5, we treated one group of mice with CNO and the other with vehicle. No significant changes in rotarod performance were observed between groups (Fig. 7h). We then kept the mice in their home cage for 7 additional days. Then, both groups of mice were treated again with vehicle or CNO on day 13 (i.e. 6^th^ day of rotarod task). On day 13 we observed a significant reduction in the accelerating rotarod performance in mice treated with CNO compared with vehicle-treated ones (Fig. 7h). To rule out the possibility of unspecific or off-target effects of CNO, we repeated exactly the same experiment as in figure 7h using a new cohort of wild type (WT) mice treated with CNO or vehicle (Fig. 7i). CNO *per se* did no induce any effect on the accelerating rotarod performance at any time or session. This result ruled out unspecific off-target effects of CNO.

We then subjected the same *Egr1*-CreER^T2^ mice transduced with hM3D(Gq) receptor to open field and novel object location tasks (Fig. 8a). The aim of these tests was to assess whether the manipulation of the *Egr1*-dependent neuronal ensemble induced during the accelerating rotarod task could affect unspecific and/or general hippocampal-dependent functions such as navigation, anxiety or spatial learning. Mice were injected 30 min before the open field test with vehicle or CNO (Fig. 8b). Mice treated with CNO did not display differences in terms of traveled distance (Fig. 8c), time spent in the center (Fig. 8d) and parallel index (Fig. 8e) during the 30 min session in the arena. In the novel object location test, mice treated with CNO showed no modification in new object location preference (Fig. 8f). These results show that the alteration of rotarod performance induced by CNO due to the activation of the *Egr1*-dependent neuronal ensemble induced during the accelerating rotarod task was specific, since general locomotor activity, anxiety, navigation and spatial learning skills were not altered.

**Figure 8.**
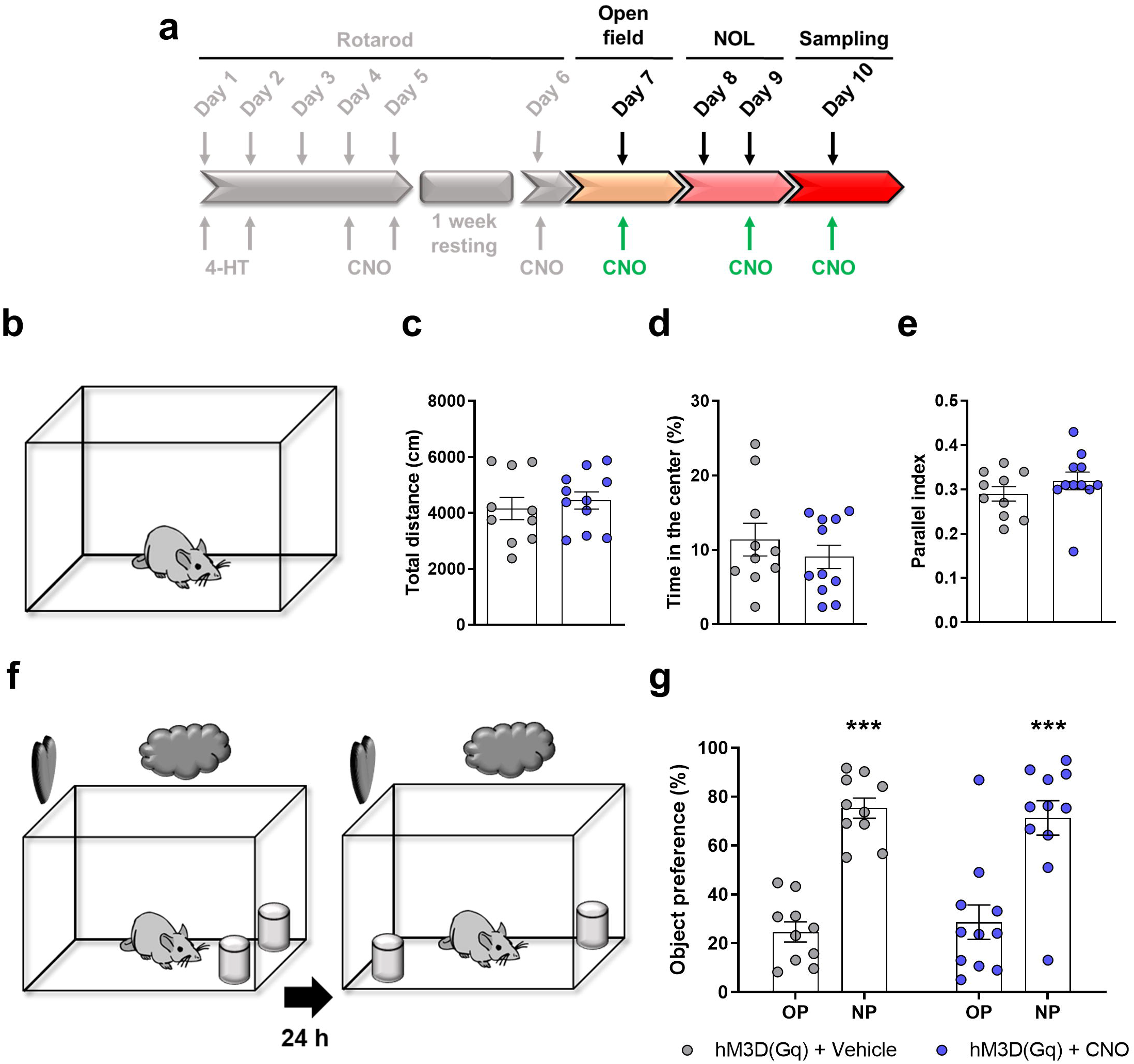
Chemogenetic modulation of the *Egr1*-dependent engrams does not alter performance in other hippocampal-related tasks. (**a**) After the accelerating rotarod task showed in figure 7 (gray tone scheme), the *Egr1*-CreER^T2^ mutant mice transduced with AAV-hSYN-DIO-hM3D(Gq)-mCherry construct and treated with 4HT at the beginning of the rotarod training (see Fig. 7) were subjected to hippocampal related tasks. Thirty min before open field task and before the testing session in the NOL task, all mice received a single i.p. injection of vehicle (n = 10) or CNO (3 mg/kg, n = 11). Additionally, on day 10, animals were treated with vehicle or CNO (3 mg/kg) 2 h before sacrifice and brains’ sampling. In the open field task (**b**), CNO injection effects on the total distance travelled (**c**), the time spent in the center of the arena (**d**) and the parallel index (**e**) are shown. Unpaired t test in **d** and **e** or Mann-Whitney t test in **g** did not detect significant differences between groups in any parameter. (**f**) In the novel object location test, (**g**) CNO injection effects on the novel object location preference are shown. Two-way ANOVA identified significant and equal novel object location preference (Novel object location preference effect: F_(1, 38)_= 62,11, *p* < 0.0001) but no differences were found between groups (Group effect: F_(1, 38)_ = 1,9 10^-14^, *p =* 0.99). OP: Old position, NP: New position. In **c**, **d**, **e**and **g** data are individual values for every mouse and means ± SEM are shown.

## Discussion

The role of specific brain regions including the striatum, motor cortex and cerebellum in the control of motor learning is well established^28–30^. More intriguing is the role of the hippocampus. Potentially, the hippocampus is capable to play either cooperative or competitive roles during the acquisition of motor skills as suggested by human studies^3,31^. However, why the hippocampus is playing such a complex role is puzzling and the underlying cellular and molecular mechanisms remain completely unknown^12^. Here we provide some clues for the first time.

Our study is the first to characterize gene regulation in the hippocampus following a motor learning task and to compare these changes to those observed in the striatum which is a core brain region regulating such behavioral skills^32^. This in-depth genome-wide regional comparative study of mRNAs showed that the acquisition of a motor skill involves common groups of genes in the striatum and hippocampus but, as expected^33–35^, unique regional transcriptional profiles can be distinguished. We identified several enriched pathways in the hippocampus related to synapse formation, modulation and reorganization, which are known to play a critical role in memories formation. Notably, when compared to non-trained mice, the hippocampus displayed major changes in gene expression in long-term trained mice whereas in short-term trained mice the changes were much more discrete. This is in line with previous reports showing that the hippocampus shows major structural changes after a long-term training procedure in the accelerating rotarod task^20^. Indeed, our cluster and pathway analysis of hippocampal transcriptomic data in response to LTT identified genes related with cytoskeleton organization, regulation of signaling transduction and transcription regulator activity which are in turn important signals for formation of hippocampal memories. The transcriptional changes observed in both regions were likely mediated by four main transcription factors (*SP1*, *Egr1*, *SMAD4* and *TEAD2*) as revealed by binding motifs analysis, suggesting that common transcriptional regulatory mechanisms are at work in the hippocampus and striatum to drive a region-specific gene regulation. Such overlapping transcriptional regulatory mechanisms have been previously reported in the amygdala and hippocampus after classical fear conditioning^36^. Importantly, *Egr1*, which is capable to induce or inhibit transcription^37,38^, is the only transcription factor that displayed changes in its expression in a task-dependent manner in the rotarod training. Our results indicate that *Egr1* is downregulated in the rotarod LTT which suggests a requirement of this down-regulation in order to permit motor learning to occur. In support of this hypothesis, knock-down of *Egr1* in CA1 improves motor learning performance.

How could *Egr1* downregulation improve motor learning? Our results provide a possible mechanism. We identify a small subset of CA1 pyramidal neurons in which *Egr1* is activated, as indicated by the Egr1/CreER^T2^ dependent labeling. This *Egr1*-dependent neuronal ensemble or engram, exerts a negative effect on motor learning or performance, as indicated by improved learning when these neurons are selectively deleted by Casp3 and impaired rotarod performance when they are chemogenetically stimulated. The simplest explanation to account for these results is that *Egr1* is induced in a subset of neurons which oppose motor learning, while general downregulation of *Egr1* in the hippocampus, favors learning, possibly by limiting the size or strength of the *Egr1*-engram. Interestingly, mice deficient for *Egr1* homolog, *Egr2*, display enhanced motor learning^39^. Thus, our results give support to a role of this CA1 neuronal ensemble as an inhibitor or controller of motor systems giving a molecular/cellular explanation of the phenomena previously described by other authors^3,31^. Whether *Egr1* could play a role in the same direction or possibly in an opposite way in the striatum is a question that needs to be further explored.

Another question in this study is, what type of information is coded by this *Egr1*-dependent neuronal ensemble during motor learning in the hippocampus? The hippocampus could play a role in the modulation of anxiety levels related to the task^40,41^, or encoding contextual information^42^ or in the regulation of goal-directed actions in synchrony with the striatum^7,12^ and even in the regulation of basal locomotion *per se*^43^. Thereby, the depletion of the *Egr1*-dependent neuronal ensemble could eliminate contextual or emotional "distracters" and, therefore, enhance the task performance. Despite the variety of information processed by the hippocampus, we did not observe compensatory changes in the striatal activation (Fig. 7e-g) or major changes in anxiety, locomotion, navigation or spatial learning processes when we manipulated the CA1 *Egr1*-dependent neuronal ensemble induced by the accelerating rotarod task (Fig. 8). Alternatively, the *Egr1*-dependent neuronal ensemble identified here could code for specific contextual information of the task. However, the hippocampus is capable to create multiple representations of the same spatial context^44^. How these representations overlap with *Egr1*-engrams and what role they may play in relation to motor learning is currently not known. In conclusion, our data show that general downregulation of *Egr1* expression levels in CA1 facilitates motor learning whereas a CA1 *Egr1*-dependent engram has a specific role in limiting motor learning. The balance between these two responses is likely to be important for optimizing behavioral responses. Thus, our study reveals a novel aspect of the role of hippocampus in the context of motor learning linked to the regulation of *Egr1*, which opens the way for further investigation of procedural memory formation.

## Methods

### Animals

For this study we used adult (12-week old) C57/BL6 males (from our colony) in experiments showed in figures 1, 2, 3 and 5. For the rest of the experiments we used the *Egr1*-CreER^T2^ mice^26^. These mice carry a bacterial artificial chromosome (BAC) including the *Egr1* gene in which the coding sequence was replaced by that of CreER^T2^ fusion protein. *Egr1*-CreER^T2^ mice were used as heterozygous in the experiments shown in figures 7 and 8 or they were crossed with R26^RCE^ mice (Gt(ROSA)26Sor^tm1.1(CAG-EGFP)Fsh^/Mmjax, Strain 004077, The Jackson Laboratory), which harbor the R26R CAG-boosted EGFP (RCE) reporter allele with a loxP-flanked STOP cassette upstream of the enhanced green fluorescent protein (EGFP) gene, to create the double heterozygous mutant *Egr1*-CreER^T2^ x R26^RCE^ mice for the experiments in figures 4 and 6. Genotypes were determined from an ear biopsy as described elsewhere^45^. For genotyping of the Cre and EGFP transgenes we used standard PCR assays following Jackson Laboratory© manufacturer’s instructions. All mice were housed together in numerical birth order in groups of mixed genotypes (3–5 mice per cage). The animals were housed with access to food and water *ad libitum* in a colony room kept at 19–22 °C and 40–60% humidity, under an inverted 12:12 h light/dark cycle (from 08:00 to 20:00). All animal procedures were approved by local committees [Universitat de Barcelona, CEEA (133/10); Generalitat de Catalunya (DAAM 5712)], in accordance with the European Communities Council Directive (86/609/EU).

### Stereotaxic surgery and viral transduction in vivo

Animals were stereotaxically injected with one of the following adeno-associated virus (AAV): AAV-flex-taCasp3-TEVp (UNC vector core); AAV-CaMKII-shRNA-*Egr1*-mCherry (#shAAV-258146, Vector Biolabs), AAV-Scramble-mCherry (#1781, Vector Biolabs), AAV-CAG-FLEX-tdTomato (UNC vector core) and pAAV-hSyn-DIO-hM3D(Gq)-mCherry (Addgene, #44361). Briefly, mice were anaesthetized with ketamine-xylazine (100 mg/kg), and bilaterally injected with AAVs (~2.6 x 10^9^ GS per injection) in the CA1 of the dorsal hippocampus, from the bregma (millimetres); antero-posterior, –2.0; lateral, ±1.5; and dorso-ventral, −1.3. AAV injection was carried out in 2 min. The needle was left in place for 7 min for complete virus diffusion before being slowly pulled out of the tissue. After 2 h of careful monitoring, mice were returned to their home cage for 3 weeks. All mice subjected to surgery that survived and were healthy without clinical problems (such as head inclination or >15% of body weight loss) were also behaviorally characterized. Once the behavioral characterization was done, half of the brain was used to verify the site of injection by immunofluorescence (see *Tissue fixation, immunofluorescence’*s section). Mice that showed no correct viral transduction were excluded from the entire study.

### MRI image acquisition and analysis

Eleven WT male mice (B6CBA background) were scanned at 18-20 weeks of age in a 7.0T BioSpec 70/30 horizontal animal scanner (Bruker BioSpin, Ettlingen, Germany), equipped with an actively shielded gradient system (400 mT/m, 12-cm inner diameter). Each animal underwent structural T2-weighted imaging, diffusion weighted imaging (DWI) and resting-state functional magnetic resonance (rs-fMRI) during the same acquisition session to evaluate connectivity between regions of interest. Animals were placed in supine position in a Plexiglas holder with a nose cone for administering anesthetic gases and were fixed using tooth an ear bars and adhesive tape. Animals were anesthetized with 2.5% isoflurane (70:30 N_2_O:O2) and a combination of medetomidine (bolus of 0.3 mg/kg, 0.6 mg/kg/h infusion) and isoflurane (0.5 %) was used to sedate the animals. 3D-localizer scans were used to ensure accurate position of the head at the isocenter of the magnet. T2-weigthed image was acquired using a RARE sequence with effective TE = 33 ms, TR = 2.3 s, RARE factor = 8, voxel size = 0.08 x 0.08 mm² and slice thickness = 0.5 mm. DWI was acquired using an EPI sequence with TR = 6 s, TE = 27.6 ms, voxel size 0.21 x 0.21 mm² and slice thickness 0.5 mm, 30 gradient directions with b-value = 1000 s/mm² and 5 baseline images (b-value = 0 s/mm²). rs-fMRI was acquired with an EPI sequence with TR = 2s, TE = 19.4, voxel size 0.21 x 0.21 mm² and slice thickness 0.5 mm. 420 volumes were acquired resulting in an acquisition time of 14 minutes.

Seed-based analysis was performed to evaluate connectivity of the left striatum to the hippocampus of both hemispheres. rs-fMRI was preprocessed, including slice timing, motion correction by spatial realignment using SPM8, correction of EPI distortion by elastic registration to the T2-weighted volume using ANTs^46^, detrend, smoothing with a full-width half maximum (FWHM) of 0.6mm, frequency filtering of the time series between 0.01 and 0.1 Hz and regression by motion parameters. All these steps were performed using NiTime (http://nipy.org/nitime). Region parcellation was registered from the T2-weighted volume to the preprocessed mean rs-fMRI. Connectivity between each pair of regions was estimated as the Fisher-z transform of the correlation between average time series in each region. Network organization was quantified using graph metrics, namely, strength, global and local efficiency and average clustering coefficient^47^. To perform the seed-based analysis, two regions-striatum and hippocampus-were selected from the automatic parcellation. For each seed region, average time series in the seed was computed and correlated with each voxel time series, resulting in correlation maps describing the connectivity of left striatum with the rest of the brain and the hippocampus was identified from brain parcellation to evaluate their connectivity with the seed region. Connectivity was quantified as the mean value of the correlation map in each region, considering only positive correlations.

### Pharmacological treatments

We used single intra-peritoneal injections of 4-hydroxytamoxifen or 4-HT (Sigma, #H7904) 50 mg/kg or clozapine-N-oxide or CNO (Sigma, #C0832) 3 mg/kg. The 4-HT’s vehicle was peanut oil (Sigma, #2144) (with a previous dissolution by heating in 100 % EtOH) and for CNO was distilled water. The 4-HT was always administered 1 h prior to the behavioral testing and the CNO was always administered 30 min prior to the behavioral testing or 2 h prior to the mice sacrifice and brain tissue collection. See schematic experimental design in each corresponding figure (1 and 5) for further details.

### Accelerating rotarod

As previously described^48^, animals were placed on a motorized rod (30-mm diameter, Panlab, Spain). The rotation speed was gradually increased from 4 to 40 r.p.m. over the course of 5 min. The fall latency time was recorded when the animal was unable to keep up with the increasing speed and fell. Rotarod training/testing was performed 4 times per day during 5 consecutive days. The results show the average of fall latencies per trial for group each day.

### Fiber photometry

Male C57BL/6 mice were anaesthetized with isoflurane and received 10 mg.kg-1 intraperitoneal injection (i.p.) of Buprécare® (buprenorphine 0.3 mg) diluted 1/100 in NaCl 9 g.L-1 and 10 mg.kg-1 of Ketofen® (ketoprofen 100 mg) diluted 1/100 in NaCl 9 g.L-1, and placed on a stereotactic frame (Model 940, David Kopf Instruments, California). 1 μL of virus (AAV9.CamKII.GCaMP6f.WPRE.SV40, titer ≥ 1×10¹³ vg/mL, working dilution 1:10), was injected unilaterally into the CA1 (L = −1.25; AP = −2; V = −1.1-1.2, in mm) at a rate of 0.1 μl.min-1. pENN.AAV.CamKII.GCaMP6f.WPRE.SV40 was a gift from James M. Wilson (Addgene viral prep # 100834-AAV9).

A chronically implantable cannula (Doric Lenses, Québec, Canada) composed of a bare optical fiber (400 μm core, 0.48 N.A.) and a fiber ferrule was implanted at the location of the viral injection site. The fiber was fixed onto the skull using dental cement (Super-Bond C&B, Sun Medical). Real time fluorescence emitted from GCaMP6f-expressing neurons was recorded using fiber photometry as described^49^. Fluorescence was collected using a single optical fiber for both delivery of excitation light streams and collection of emitted fluorescence. The fiber photometry setup used 2 light-emitting LEDs: 405 nm LED sinusoidally modulated at 330 Hz and a 465 nm LED sinusoidally modulated at 533 Hz (Doric Lenses) merged in a FMC4 MiniCube (Doric Lenses) that combines the 2 wavelengths excitation light streams and separate them from the emission light. The MiniCube was connected to a Fiberoptic rotary joint (Doric Lenses) connected to the cannula. A RZ5P lock-in digital processor controlled by the Synapse software (Tucker-Davis Technologies, TDT, USA), commanded the voltage signal sent to the emitting LEDs via the LED driver (Doric Lenses). The light power before entering the implanted cannula was measured with a power meter (PM100USB, Thorlabs) before the beginning of each recording session. The irradiance was ~9 mW/cm2. The fluorescence emitted by GCaMP6f in response to light excitation was collected by a femtowatt photoreceiver module (Doric Lenses) through the same fiber patch cord. The signal was received by the RZ5P processor (TDT). Real time fluorescence due to 405-nm and 465-nm excitations was demodulated online by the Synapse software (TDT). A camera was synchronized with the recording using the Synapse software. Signals were exported to MATLAB R2016b (Mathworks) and analyzed offline. After careful visual examination of all trials, they were clean of artifacts in these time intervals. The timing of events was extracted from the video.

To calculate ∆F/F, a linear least-squares fit was applied to the 405 nm signal to align it to the 465 nm signal, producing a fitted 405 nm signal. This was then used to normalize the 465 nm signal as follows: ∆F/F = (465 nm signal-fitted 405 nm signal)/fitted 405 nm signal^50^. For each trial, signal analysis was performed for 2 time windows: from −10 to +20 sec around the moment the mouse is positioned on the rotarod (onset) and from −10 sec before the fall of the animal to the end of the trial (fall). The percentage of change of the AUC was calculated between [0 10] and [10 20]sec after onset and between [−10 −5] and [−5 0] before fall.

### Open field and novel object location test

For the novel object location test (NOL), an open-top arena (45 × 45 × 45 cm) with visual cues surrounding the apparatus was used. Mice were first habituated to the arena (1 day, 30 min). We considered this first exposition to the open arena as an open field paradigm. We monitored total traveled distance, time spent in the center of the arena and parallel index as measures of locomotor activity, anxiogenic behavior and spatial navigation strategies respectively. On day 2, two identical objects (A1 and A2) were placed in the arena and explored for 10 min. Twenty-four hour later (Day 3), one object was moved from its original location to the diagonally opposite corner and mice were allowed to explore the arena for 5 min. The object preference was measured as the time exploring each object × 100/time exploring both objects. Behavioral data was processed and analyzed using the Smart Junior software (Panlab, Spain).

### RNA sequencing analysis

#### RNA Extraction and Quality Control

Hippocampal and striatal samples were homogenized, and RNA extracted using RNeasy Lipid Tissue Mini kit (Quiagen) according to manufacturer’s recommendations. RNA purity and quantity were determined with a UV/V spectrophotometer (Nanodrop 1000), while RNA integrity was assessed with a 2100 Bioanalyzer (Agilent Technologies Inc., CA), according to manufacturers’ protocols. The average RIN value for our samples was 9.5, and the RIN cut-off for sample inclusion was 8.0.

#### RNA Sequencing and Differential Gene Expression Analysis

Libraries were prepared using the TruSeq Stranded mRNA Sample Prep Kit v2 (ref. RS-122-2101/2) according to the manufacturer’s protocol. Briefly, 500 ng of total RNA were used for poly(A)-mRNA selection using streptavidin-coated magnetic beads and were subsequently fragmented to approximately 300 bp. cDNA was synthesized using reverse transcriptase (SuperScript II, ref. 18064-014, Invitrogen) and random primers. The second strand of the cDNA incorporated dUTP in place of dTTP. Double-stranded DNA was further used for library preparation. dsDNA was subjected to A-tailing and ligation of the barcoded Truseq adapters. Library amplification was performed by PCR using the primer cocktail supplied in the kit. All purification steps were performed using AMPure XP beads. Final libraries were analyzed using Fragment Analyzer to estimate the quantity and check size distribution, and were then quantified by qPCR using the KAPA Library Quantification Kit (ref. KK4835, KapaBiosystems) prior to amplification with Illumina’s cBot. Sequencing was done using the HiSeq2500 equipment (illumina), Single Read, 50bp, using the v4 chemistry.

The quality of the sequencing data was checked using the FastQC software v0.11.5. Andrews S. (2010). FastQC: a quality control tool for high throughput sequence data. Available online at: http://www.bioinformatics.babraham.ac.uk/projects/fastqc.

An estimation of ribosomal RNA in the raw data was obtained using riboPicker version 0.4.3^51^. Reads were aligned to the GENCODE version of the *Mus musculus* genome, release M20 (GRMm38/mm10 assembly) using the STAR mapper (version 2.5.3a)^52^. The raw read counts per gene was also obtained using STAR (--quantMode TranscriptomeSAM GeneCounts option) and the GENCODE release M20 annotation (ftp://ftp.ebi.ac.uk/pub/databases/gencode/Gencode_mouse/release_M20/gencode.vM20.annotation.gtf.gz). The R/Bioconductor package DESeq2 version.22.2 (R version 3.5.0) was used to assess the differentially expressed genes between experimental groups, using the Wald statistical test and the False Discovery Rate for the p-value correction. Prior to the differential expression analysis, genes with the sum of raw counts across all samples below 10 were discarded, the library sizes were normalized using the default DeSeq2 method, and the read counts were log2 transformed. To exclude false positive genes, genes with low expression levels (baseMean <10) were excluded from the list of DEGs.Sequencing data has been deposited in NCBI’s Gene Expression Omnibus and are accessible through GEO Series accession number (Accession number pending).

#### Gene Functional Enrichment Analysis

Panther pathway enrichment analysis was performed to explore the functional roles of DEGs in the paired comparison of striatum and hippocampus mRNA-seq data. To discover further possible connections between DEGs and transcription factors we used EnrichR. This web-based tool which computes enrichment through 35 different gene-set libraries. We detected the binding motif sites in our gene list using the position weight matrices (PWMs) analysis from TRANSFAC and JASPAR. The PWMs from TRANSFAC and JASPAR were used to scan the promoters of all mouse genes in the region between −2,000 and +500 from the transcription factor start site (TSS). Functional annotations for modules of interest were generated using the web server SynGO (https://www.syngoportal.org/), which provides an expert-curated resource for synapse function and gene enrichment analysis^55^.

#### Reverse Transcription and Real-time qPCR

For gene expression analysis, 500 ng of total RNA were retrotranscribed using the High Capacity cDNA Reverse Transcription Kit (Applied Biosystems, ref. 4368814) according to manufacturer’s instructions. The resulting cDNA was analyzed by qPCR using the following gene expression assays (PrimeTime qPCR Assays, Integrated DNA Technologies, Inc.): Egr-1 (Mm.PT.58.29064929) and Actinβ (Mm.PT.39a.22214843.g). The reaction was performed in a final volume of 12 μL using the Premix Ex Taq Probe based qPCR assay (Takara Biotechnology, ref. RR390A). All reactions were run in duplicate. Data analysis was performed using the MxProTM qPCR analysis software version 3.0 (Stratagene). Relative enrichment was calculated using the ΔΔCt method, with Actinβ serving as an internal loading control.

### Tissue fixation, immunofluorescence and confocal imaging

Animals were deeply anaesthetized and subsequently intracardially perfused with 4% (weight/vol) paraformaldehyde in 0.1□M phosphate buffer. The brains were dissected out and kept 48 h in 4% paraformaldehyde. Sagittal sections (40□μm) were obtained using a vibratome (Leica VT1000). For immunofluorescence, after blocking/permeabilization (1□h in PBS containing 3 mL/L Triton X-100 and 10 g/L bovine serum albumin (BSA)), sections were incubated overnight with specific antibodies against MAP2 (1:500; #M1406, Sigma-Aldrich, St. Louis, MO, USA), DARPP32 (1:500, #611520, BD Transductions), NeuN (1:500, #MAB377, Millipore/Chemicon), Parvalbumin (1:1000, #PV27, SWANT), GFP FITC-conjugated (1:500, #Ab6662, Abcam), Egr1 (1:1000, #4154S, Cell Signalling) and cFos (1:150, #sc-52, Santa Cruz Biotechnology). After incubation (2 h) with appropriate fluorescent secondary antibodies (Cy3-or Cy2-coupled fluorescent secondary antibodies, 1:200; Jackson ImmunoResearch Laboratories catalog #715-165-150 and #715-545-150 respectively), nuclei were stained (10 min) with 4′,6-diamidino-2-phenylindole (DAPI; catalog #D9542, Sigma-Aldrich). The sections were mounted onto gelatinized slides and cover-slipped with Mowiol.

### Image analysis and pseudo-stereological counting

Images (at 1024 × 1024 pixel resolution) in a mosaic format were acquired with a Leica Confocal SP5 with a□×□40 oil-immersion or x20 normal objectives and standard (1 Airy disc) pinhole (1 AU) and frame averaging (3 frames per *z* step) were held constant throughout the study. For pseudo-stereological counting, we analyzed 3 sagittal sections, from 1.4 to 2.0 mm relative to bregma (Supplementary figure 1), spaced 300□μm apart. The areas of analysis were, dorsal CA1, dorsal CA3, dorsal striatum, ventral striatum and layers 2/3, 4, 5 and 6 of the motor cortex area 1 (Supplementary figure 1). Unbiased blind counting of GFP-or MAP2-or DARPP32-or Parvalbumin-or NeuN-positive neural cells relative to genotype and condition was performed and normalized to the area of counting.

### Statistics

Analyses were done using Prism version 6.00 for Windows (GraphPad Software, La Jolla, CA, USA). Data are expressed as means□± SEM Normal distribution was tested with the d’Agostino and Pearson omnibus test. If no difference from normality was detected, statistical analysis was performed using two-tailed Student’s *t* test or ANOVA and Tukey’s or Dunnett’s post-hoc tests. If distribution was not normal, non-parametric two-tailed Mann-Whitney test was used. The *p*□<□0.05 was considered as significant. All statistical row data are summarized in supplementary table 7.

## Acknowledgements

AG is a Ramón y Cajal fellow (RYC-2016-19466) supported by a grant from Ministerio de Ciencia, Innovación y Universidades (RTI2018-094678-A-I00). We thank Ana López (María de Maeztu Unit of Excellence, Institute of Neurosciences, University of Barcelona, MDM-2017-0729, Ministry of Science, Innovation and Universities) for technical support. We thank María Calvo from the Advanced Microscopy Service (Centres Científics i Tecnològics Universitat de Barcelona) for her help in the acquisition, analysis and interpretation of the confocal images. We also thank to the Experimental MRI 7T Unit of the IDIBAPS and to CERCA Programme (Generalitat de Catalunya) and to Guadalupe Sòria, Emma Muñoz-Moreno and Javier López-Gil for their technical support. We also thank to Daniel del Toro and Eulàlia Martí, from the Institut de Neurociències, for their insightful comments and advices.

## Author contributions

V.B. conceived and carried out most experiments, analyzed and interpreted results and wrote the manuscript. E.M. carried out experiments related to fiber photometry. A.P. performed and took part in the mRNA and RNAseq experiments. M.M. designed, analyzed and interpreted the fMRI experiment. S.G. supervised the behavioral experiments. J.A. discussed and supervised histological and morphological experiments. C.M. supervised and designed fiber photometry experiments. J-A.G. supervised and interpreted neuronal ensembles formation experiments. A.G. conceived and supervised the study, analysed results and wrote the manuscript.

---

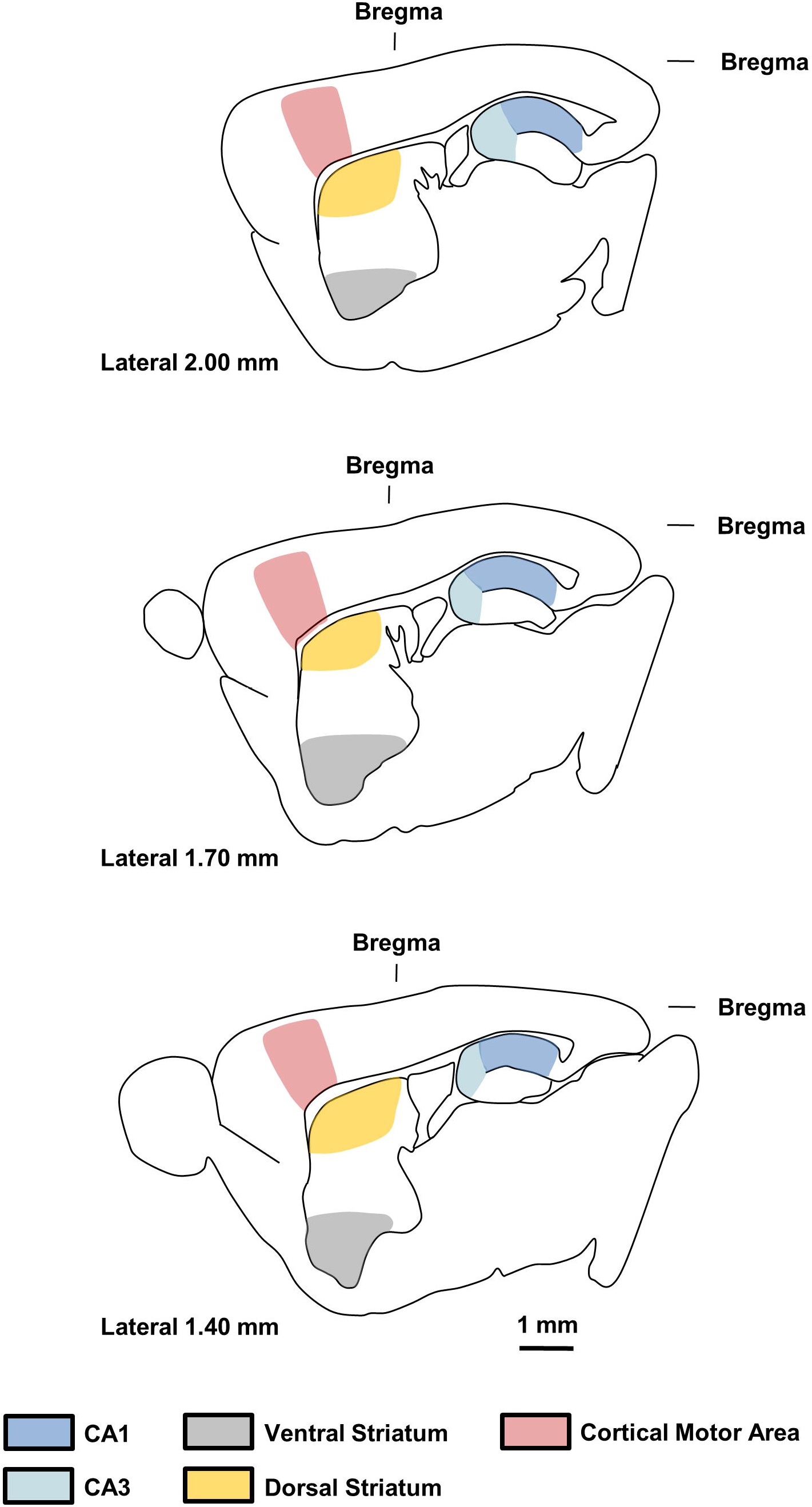

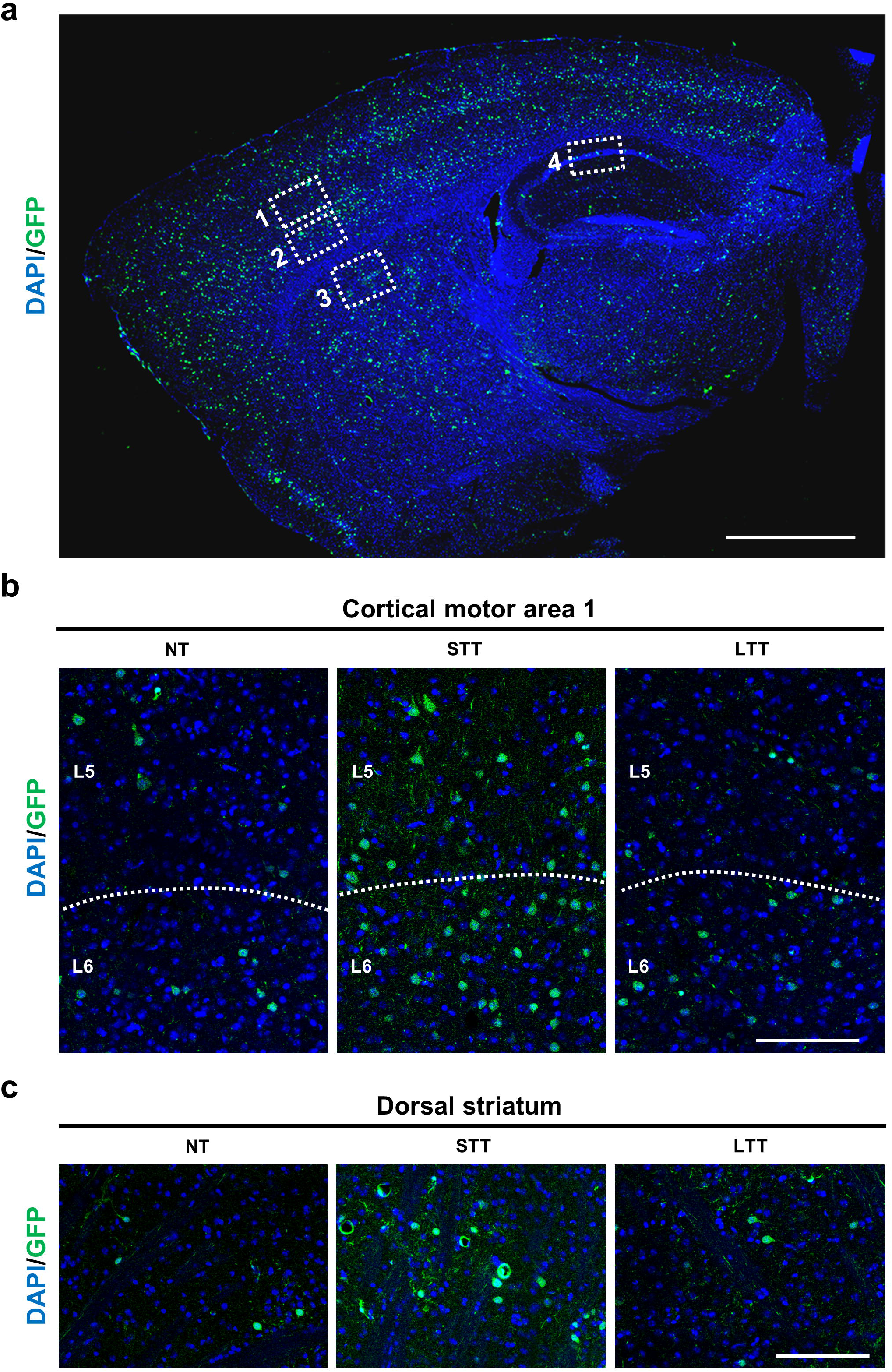

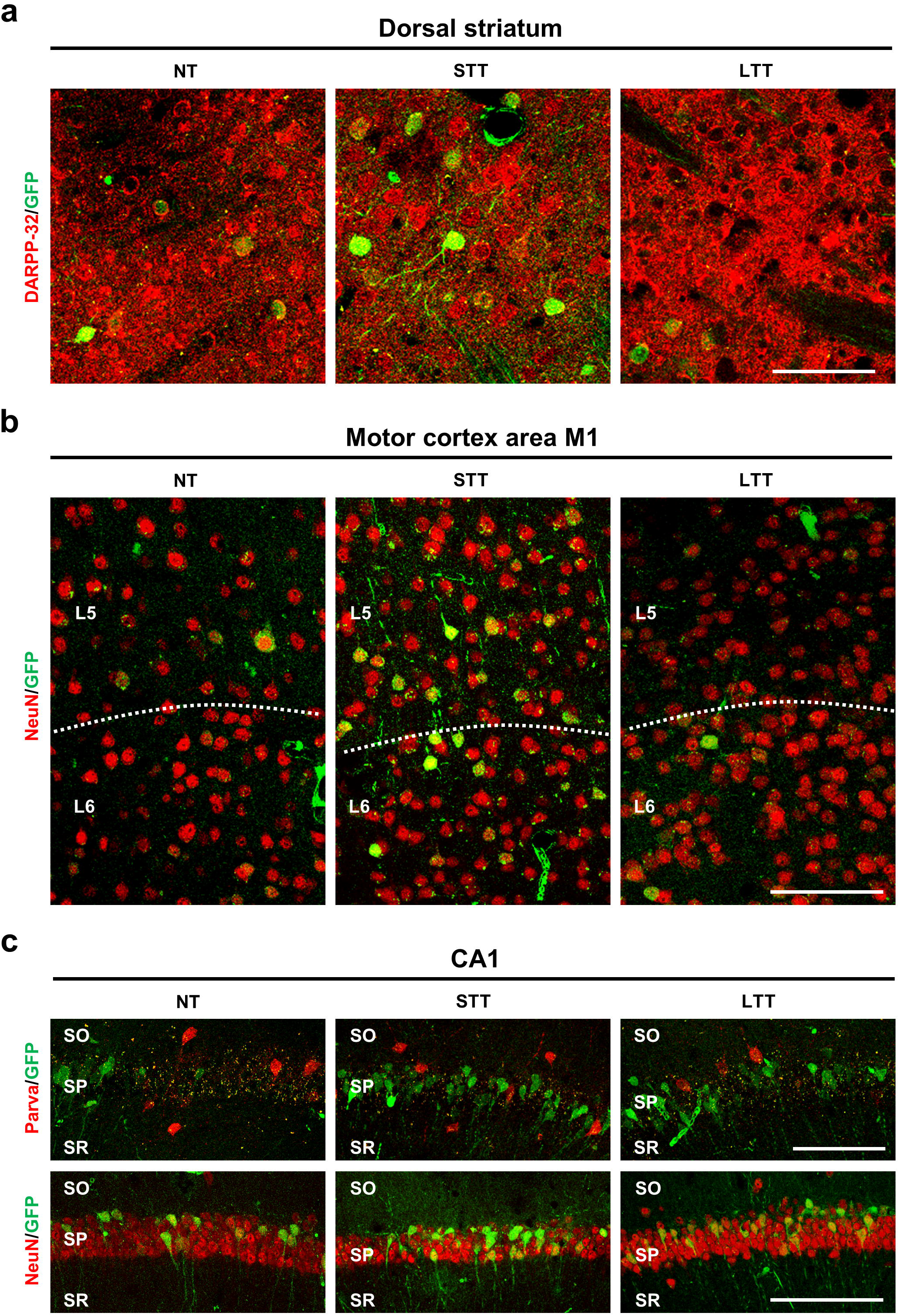

