## Supplementary figure legends for "*Hippocampal Egr1*-dependent neuronal ensembles negatively regulate motor learning"

**Supplementary figure 1. Schematic representation of the brain regions evaluated in the Egr1-CreERT2 x RCE:LoxP mice after the accelerating rotarod test performance.** (**a**) Representative schematic illustration of the three slices per mouse evaluated in figure 3. Sagittal sections were evaluated. Slices were taken 300 microns apart from 2.00 mm to 1.40 mm respect to bregma. Scale bar: 1 mm.

**Supplementary figure 2. Representative brain areas with significant differences in the Egr1-CreERT2 x RCE:LoxP mice subjected to the accelerating rotarod task.** (**a**) Representative image of Egr1-dependent activation of neural cells (GFP-positive, green) co-stained with DAPI (blue) in an entire sagittal section from an Egr1-CreERT2 x RCE:LoxP mouse. White rectangles depict brain sub-regions with significant differences between groups; 1: Layer 5 of the motor cortical area, 2: Layer 6 or the motor cortical area, 3: Dorsal striatum and 4: Hippocampal CA1. (**b**) Representative insets of layers 5 and 6 of the cortical motor area in the three groups of Egr1-CreERT2 x RCE:LoxP mice: Non-trained mice (NT), short-term trained mice (STT) and long-term trained mice (LTT) in the accelerating rotarod task. (**c**) Representative insets of the dorsal striatum in the three groups of Egr1-CreERT2 x RCE:LoxP mice: Non-trained mice (NT), short-term trained mice (STT) and long-term trained mice (LTT) in the accelerating rotarod task. Scale bar in A: 1 mm. Scale bar in B and C: 70 microns.

**Supplementary figure 3. Representative images of neuronal subpopulations activated in brain regions of Egr1-CreERT2 x RCE:LoxP mice trained in the accelerating rotarod task.** (**a**) Representative images of Egr1-dependent activation of neural cells (GFP-positive, green) co-stained with DARPP-32 in the striatum (**b**), NeuN in the motor cortex area (**c**) and with Parvalbumin and MAP2 in the hippocampal CA1 (**d**). DARPP-32, NeuN, Parvalbumin and MAP2 are depicted in red. All sections are from Egr1-CreERT2 x RCE:LoxP mice subjected to the three conditions: Non-trained mice (NT), short-term trained mice (STT) and long-term trained mice (LTT) in the accelerating rotarod task. Scale bar in A: 80 microns. Scale bar in B and C: 100 microns. L5: Layer 5 of the motor cortex area, L6: Layer 6 of the motor cortex area, CA1: Cornu amonis 1, SO: *Stratum oriens*, SP: *Stratum pyramidale*, SR: *Stratum radiatum*.
